# ScIsoX: A Multidimensional Framework for Measuring Isoform-Level Transcriptomic Complexity in Single Cells

**DOI:** 10.1101/2025.04.28.650897

**Authors:** Siyuan Wu, Ulf Schmitz

## Abstract

Single-cell isoform sequencing enables high-resolution characterisation of transcript isoform expression, yet analytical frameworks to systematically measure transcriptomic complexity are lacking. Here, we introduce `ScIsoX`, a computational framework that integrates a novel hierarchical data structure, a suite of complexity metrics, and dedicated visualisation tools for isoform-level analysis. `ScIsoX` supports systematic exploration of global and cell-type-specific isoform expression patterns arising from alternative splicing, revealing multidimensional complexity signatures across diverse datasets - insights often missed by conventional gene-level approaches.

---

Alternative splicing dramatically expands the functional repertoire of eukaryotic cells by generating diverse transcript isoforms from a limited number of genes. Recent advances in single-cell long-read sequencing (SCLRS) have enabled comprehensive profiling of full-length transcript isoforms at unprecedented resolution [1]. However, analytical frameworks for measuring and interpreting the multidimensional nature of transcriptomic complexity at single-cell resolution do not exist. This represents a missed opportunity to leverage the additional layers of information provided by SCLRS data, which this study aims to address.

Current approaches for analysing SCLRS data face two major challenges. First, conventional data structures, such as gene-by-cell count matrices, inherently fail to capture the complexity and variability of isoform usage across genes. Attempts to merge gene-level and isoform-level count matrices into a “cell × gene × isoform” tensor necessitate extensive zero padding to accommodate gene-specific variability in isoform numbers, resulting in sparse 3D tensors with excessive memory demands. Second, while existing analytical methods excel at isoform discovery and quantification [2, 3], they lack comprehensive metrics that address fundamental questions about the organising principles governing isoform expression patterns across cells and cell types.

To address these challenges, we introduce `ScIsoX`, a computational framework that implements (i) a novel Single-Cell Hierarchical Tensor (SCHT) data structure, (ii) a comprehensive suite of analytical metrics, and (iii) visualisation tools for measuring transcriptomic complexity across multiple biological scales (Fig.1a and Supplementary Fig.1). At its core, the SCHT organises isoform-level count data into gene-specific sub-tensors, where each gene is represented by an individual count matrix containing isoform-by-cell expression values. This partition-based design preserves the intrinsic hierarchy without resorting to extensive zero padding, yielding a representation that is both biologically meaningful and computationally efficient. When cell type information is integrated, the SCHT is extended to include cell types as an additional dimension. Each count matrix contains only the cells belonging to that particular cell type expressing the gene, creating a multi-level hierarchy that elegantly captures gene-isoform-cell relationships.

**Figure 1:**
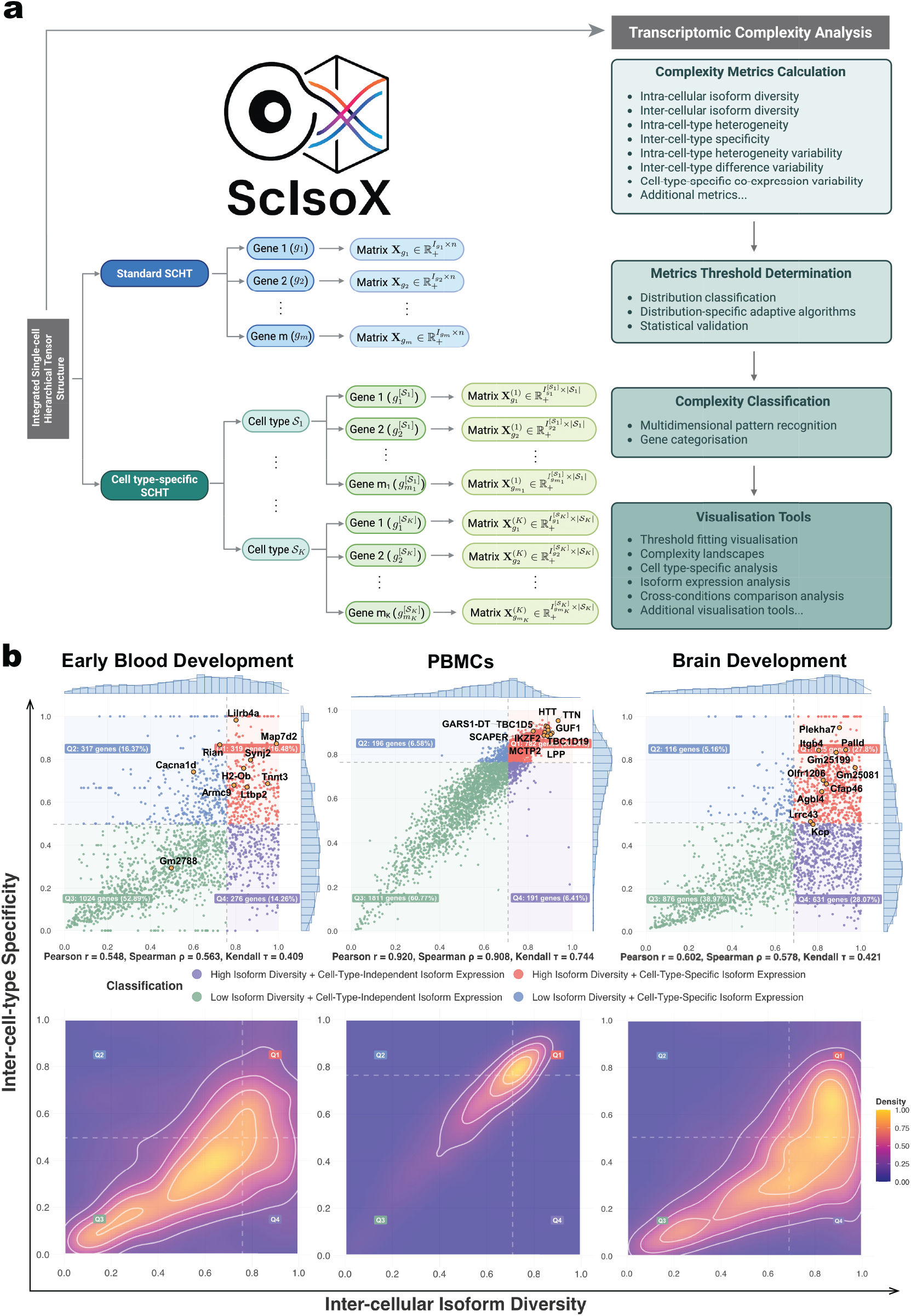
ScIsoX overview and comparison of complexity landscapes across biological systems. (a) Core of the `ScIsoX` computational framework showing the interconnected components of data structure and analytical processing. Left: SCHT construction organises isoform expression data into gene-specific sub-tensors; Right: `ScIsoX`’s analytical pipeline progressing from complexity metrics to biological insights. Created with BioRender.com. (b) Complexity landscapes in mouse early blood development, human peripheral blood mononuclear cells, and mouse brain development datasets, illustrating the relationship between selected complexity dimensions. Top: The visualised complexity space reveals gene distribution across four quadrants, with annotated genes of interest that demonstrate characteristic complexity signatures; Bottom: Density contour maps revealing system-specific clustering patterns, demonstrating how complexity distributions vary across different biological contexts.

Building upon this structure, `ScIsoX` conceptualises transcriptomic complexity through seven core metrics, each capturing a distinct dimension of isoform expression patterns (Fig.1a and Supplementary Table 1). The primary dimensions include (I) intra-cellular isoform diversity (i.e., the tendency for a gene to co-express multiple isoforms within individual cells), (II) inter-cellular isoform diversity (i.e., the diversity of isoforms expressed by a gene across the whole cell population), (III) intra-cell-type heterogeneity (i.e., cell-to-cell variation in isoform usage), and (IV) inter-cell-type specificity (i.e., measure of cell-type-specific isoform usage). Three additional higher-order metrics measure variability in these patterns to determine, (V) whether cellular heterogeneity is concentrated in specific cell types, (VI) whether cell-type-specific differences occur between particular lineages, and (VII) whether isoform co-expression patterns vary across cell types. To complement these core metrics, we provide additional characterisation metrics that capture specific aspects of isoform usage (Supplementary Table 2).

We have confirmed `ScIsoX`’s utility by analysing three distinct SCLRS datasets surveying: (1) murine haematopoietic development via Nanopore sequencing [4], (2) murine and human brain development via Nanopore sequencing [5], and (3) human peripheral blood mononuclear cells (PBMCs) via PacBio’s Kinnex protocol [6]. These datasets represent fundamentally different biological systems while also employing distinct technical approaches to long-read sequencing. This selection enabled comprehensive evaluation of our framework’s performance and broad applicability. All datasets included robust cell type annotations for analysis. Our analysis revealed markedly different transcriptomic complexity patterns in these systems, highlighting the biological insights uniquely accessible through our approach.

The transcriptomic complexity analysis implemented in `ScIsoX` can, for example, assess distinct isoform expression patterns (Fig.1b). These patterns were non-randomly distributed, with murine haematopoietic development exhibiting a bimodal pattern dominated by low isoform diversity and low cell type specificity (Q3: 52.89%) with fewer genes showing cell-type-specific expression (Q1+Q2: 32.85%) (Fig.1b). The mouse brain development dataset exhibited a similar bimodal pattern, also demonstrating substantial diversity across quadrants with notable clusters. In contrast, the human PBMC dataset exhibited a strikingly different distribution compared to the two development datasets, showing a remarkably strong positive correlation between inter-cellular isoform diversity and inter-cell-type specificity (Fig.1b). This tight correlation suggests that in specialised immune cells, isoform diversity is closely linked to cell-type-specific functions. Both developmental datasets showed a greater range of specificity/diversity relationships than PBMCs, reflecting greater transcriptomic heterogeneity in development compared to specialised immune cells, which require specific isoform-switching events for state transitions and to respond to cellular signals. Our framework uniquely identifies genes with interesting complexity profiles that may be overlooked by conventional SCLRS analysis. For instance, the vast majority of genes in all datasets exhibit higher inter-cellular diversity compared to intra-cellular diversity, demonstrating a fundamental principle: genes tend to express cell-type-specific isoforms rather than multiple isoforms in each cell type (Fig.2a). However, a subset of genes with intra-cellular diversity that is higher than their inter-cellular diversity can be identified, suggesting coordinated co-expression of multiple isoforms within individual cells rather than cell-specific isoform selection. These genes may require specific interdependent isoform relationships for proper function, representing a distinct regulatory mechanism for further study. For example, while the role of *Sox17* in endothelial-to-haematopoietic transition is well-established [7, 8], the specific significance of its multiple transcript isoforms remains largely unexplored. Our analysis suggests that *Sox17* may utilise coordinated expression of multiple isoforms to achieve its diverse regulatory functions during early haematopoietic development (Supplementary Fig.2).

**Figure 2:**
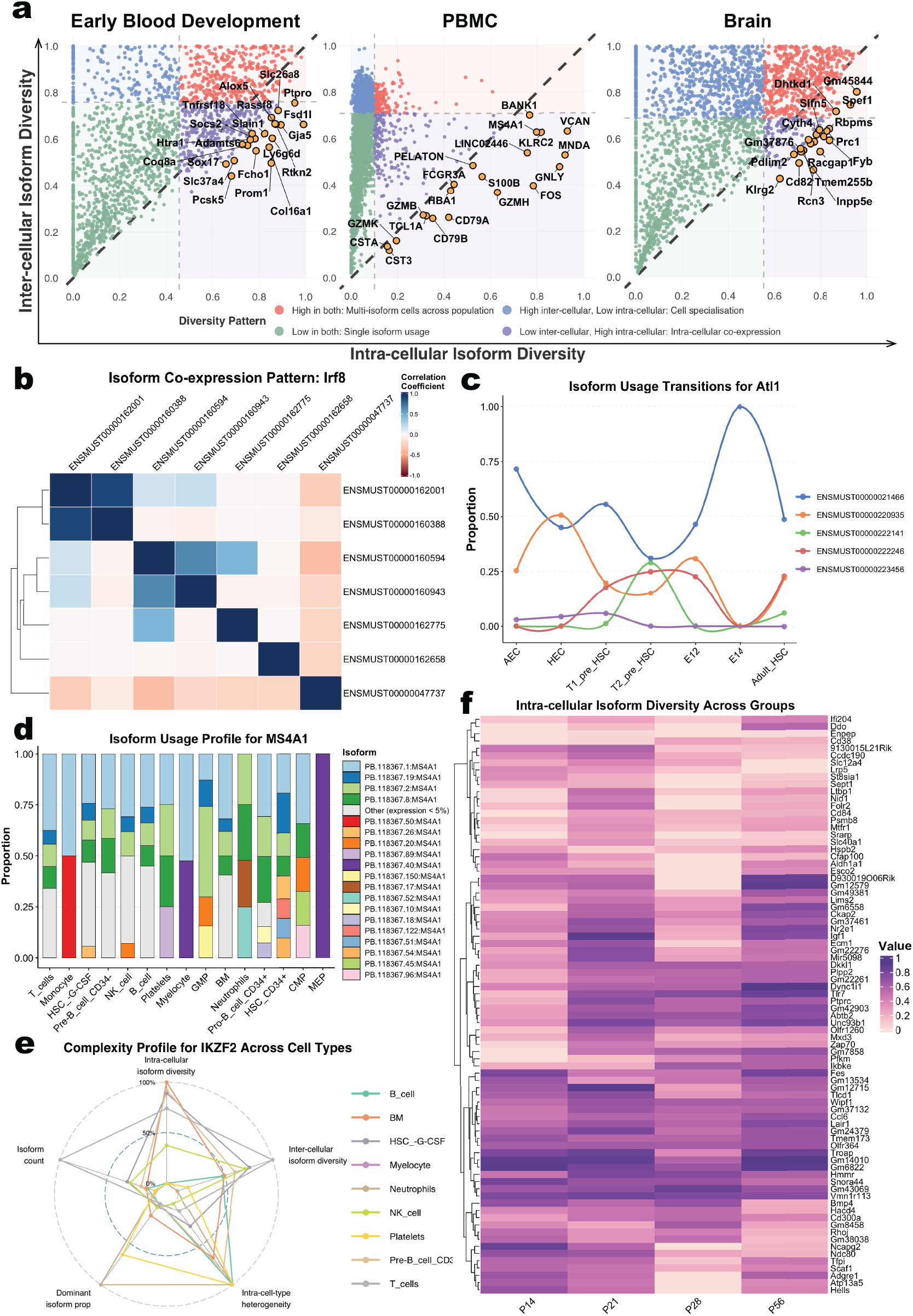
Multidimensional transcriptomic complexity analysis reveals isoform expression patterns. (a) Intra-cellular versus inter-cellular diversity analysis across three datasets. Highlighted genes falling below the diagonal line (i.e., where intra-cellular diversity exceeds inter-cellular diversity). (b) *Irf8* isoform co-expression correlation analysis, showing both positive and negative expression correlations between different isoforms, suggesting complex regulatory relationships. (c) *Alt1* isoform proportion transitions during mouse haematopoietic development. (d) *MS4A1* isoform usage profiles across different immune cell types in PBMCs. (e) Comparison of *IKZF2* complexity profiles across different immune cell types in PBMCs. (f) Heatmap of intra-cellular isoform diversity across brain postnatal developmental stages (Days 14, 21, 28 and 56).

Co-expression analysis reveals distinct patterns of coordinated isoform expression. For instance, in murine haematopoietic development, the transcription factor *Irf8*, a key interferon regulatory factor critical for myeloid lineage determination and immune cell differentiation [9], shows multiple clusters of co-expressed isoforms (Fig.2b). `ScIsoX` also enables tracking proportions of expressed isoforms across cell types, further highlighted dynamic changes in isoform usage, e.g., across lineages or developmental stages (Fig.2c). In addition, `ScIsoX` facilitates detailed examination of genes’ cell-type-specific complexity profiles. For example, the B-lymphocyte antigen *CD20* (encoded by *MS4A1*) exhibits distinctive isoform expression patterns across human PBMCs, with related myeloid cell types like myelocytes and neutrophils showing markedly different isoform preferences (Fig.2d), suggesting specialised isoform-mediated functions across immune cell lineages. Notably, *MS4A1* falls below the diagonal in the diversity analysis (Fig.2a), suggesting its function depends on the orchestrated interplay of multiple isoforms across diverse immune cell types.

Unlike existing approaches that treat isoform diversity as a single dimension, `ScIsoX` provides both a multifaceted view of transcriptomic complexity (Fig.2e and Supplementary Fig.3 - 4) and enables researchers to generate testable hypotheses about the functional significance of alternative splicing, such as across developmental timepoints or anatomical regions. For example, `ScIsoX` reveals distinct patterns of intra-cellular isoform diversity across postnatal developmental stages, with clear gene clusters exhibiting stage-specific isoform expression profiles. The heatmap in Fig.2f illustrates how certain gene groups maintain consistently high diversity (dark purple) throughout development, while others show stage-specific diversity patterns. Additionally, `ScIsoX` reveals distinct patterns of inter-cellular isoform diversity and inter-cell-type specificity that evolve dynamically throughout brain development and differ markedly between brain regions (Supplementary Fig.5 - 6).

In summary, `ScIsoX` establishes the first comprehensive framework for measuring and visualising isoform-level transcriptomic complexity in single-cell sequencing data. Through its hierarchical data structure, `ScIsoX` captures distinct dimensions of complexity at the gene, cell type, and cell population levels, generating isoform-level insights into transcriptome regulation often missed by conventional gene-level analyses. `ScIsoX`’s complexity metrics and intuitive visualisations provide a foundation for investigating the functional roles of alternative splicing in cellular differentiation, cell type specificity, and disease contexts across diverse biological systems. By using standard R objects for its core data structures and metrics, `ScIsoX` creates opportunities for future integration with other omics layers and analytical methods. Notably, while `ScIsoX` was used with SCLRS data in this study, it also processes isoform count matrices derived from single-cell short-read technologies via tools like `Kallisto-bustools` [10]. This compatibility extends its multidimensional complexity framework to existing short-read datasets, although users should consider that inherent limitations of short-reads in resolving full-length transcripts may affect certain complexity metrics.

## Methods

### Single-cell hierarchical tensor creation

`ScIsoX` introduces a novel hierarchical data structure that efficiently represents the three-dimensional relationship between genes, isoforms, and cells. Unlike conventional approaches that use either separate gene/transcript matrices, our approach organises isoform-level data into gene-specific sub-tensors. Each gene is represented by an individual matrix containing isoform-by-cell expression values, preserving the intrinsic hierarchy without extensive zero padding. While we refer to our data structure as a “hierarchical tensor”, we intentionally diverge from the strict mathematical definition, instead adopting a biologically-oriented representation specifically tailored to SCLRS data. This data structure emphasises functional utility and simplicity while facilitating scalable analysis at the isoform level to directly confront the intricacy of transcriptomic complexity.

### Quality control and normalisation

Quality control and normalisation are performed in `ScIsoX`. It requires the following inputs: (i) a raw gene count matrix, (ii) a raw isoform count matrix, (iii) a transcript annotation file, and (iv) cell metadata (optional). For the example datasets, genes were filtered that were detected in fewer than *p*_min_ proportion of cells (default: 0.02) and with mean expression counts below *ε* (default: 1 × 10^−4^). All transcripts belonging to retained genes were kept to preserve complete isoform diversity information. At the cell level, we employed a data-adaptive approach to identify and exclude low-quality cells and potential doublets based on the distribution of detected genes using the `plot_genes_per_cell_distribution()` and `recommend_qc_parameters()` functions. Cells with fewer than *n*_min_ genes (default: 200) or more than *n*_max_ genes (default: 10,000) were excluded.

Retained count data were normalised using counts per million (CPM) with subsequent logarithmic transformation:

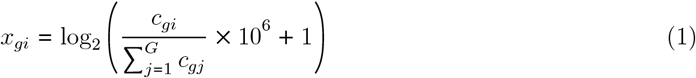

where *c*_*gi*_ represents the raw count for feature *g* in cell *i*, and *G* being the total number of features (genes or transcripts).

### Identification of highly variable genes

To prioritise computational resources on genes exhibiting biologically meaningful variation, we implemented a dispersion-based selection of highly variable genes (HVGs). For each gene *g*, `ScIsoX` calculates the variance-to-mean ratio based on their normalised expression counts. Genes are ranked by their dispersion values, and the top *n*_HVG_ genes (default: 3,000) are selected for subsequent analysis. This approach effectively identifies genes with significant biological variability whilst excluding stably expressed housekeeping genes and technical noise. If a greater number of genes is desired for inclusion, *n*_HVG_ can be increased up to the total number of genes in the dataset.

### Mathematical formulation of the SCHT data structure

We developed a hierarchical structure to represent SCLRS data. Let 𝒞 = {*c*_1_, *c*_2_, …, *c*_*n*_} be the set of all cells after quality control and filtering, and *𝒢* = {*g*_1_, *g*_2_, …, *g*_*m*_} be the set of HVGs. For each HVG *g* ∈ *𝒢* with *I*_*g*_ isoforms measured across *n* cells, we define a gene-specific expression matrix:

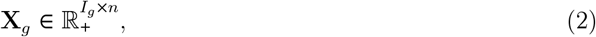

where each element *x*_*ij*_ ∈ **X**_*g*_ represents the normalised expression of isoform *i* of gene *g* in cell *j*. Note that *I*_*g*_ represents the total number of isoforms for gene *g*, and different genes may have different numbers of isoforms. The standard SCHT data structure is defined as the collection *𝒯* = {(*g*, **X**_*g*_) ∣ *g* ∈ 𝒢}. Cell-type-specific sub-tensors are created when cell type information is available. We partition the filtered cell set 𝒞 into *K* non-overlapping cell types 𝒮 = {𝒮_1_, *𝒮*_2_, …, *𝒮*_*K*_}. Note that not all genes are expressed in every cell type. For each HVG 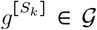 that is expressed in cell type 𝒮_*k*_ ∈ 𝒮, where *k* = 1, 2, ⋯, *K*, we denote 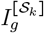 as the number of isoforms of gene *g* that are expressed in cell type 𝒮_*k*_. We then define a cell-type-specific expression matrix 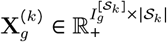 containing columns corresponding to cells of type 𝒮_*k*_. The integrated SCHT data structure with cell-type-specific structure is then defined as follows,

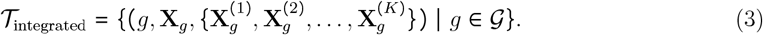

This hierarchical representation facilitates comprehensive analysis of transcriptomic complexity.

### Multi-dimensional transcriptomic complexity framework

We developed a comprehensive transcriptomic complexity analysis framework that quantifies key aspects of transcriptomic complexity, focusing on a core set of metrics that capture the essential dimensions of isoform expression patterns (Supplementary Table 1).

#### I: Intra-cellular isoform diversity

To quantify isoform diversity within individual cells, `ScIsoX` computes a weighted Shannon entropy measure for each gene. For cell *c*_*j*_ ∈ *𝒞* expressing gene *g* ∈ *𝒢*, the normalised Shannon entropy is defined as

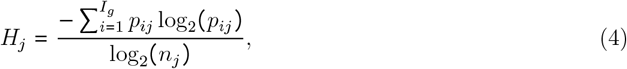

where 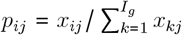 represents the proportion of gene expression attributed to isoform *i* in cell *c*_*j*_, and *n*_*j*_ is the number of isoforms detected in that cell. To account for expression magnitude, we compute a weighted mean across cells as intra-celluar isoform diversity,

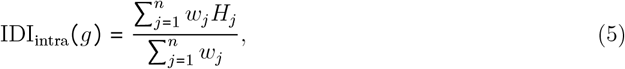

where 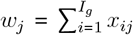 is the total expression of gene *g* in cell *c*_*j*_. This metric measures the tendency for genes to co-express multiple isoforms within individual cells.

#### II: Inter-cellular isoform diversity

To assess isoform diversity at the cell population level, `ScIsoX` computes the Shannon entropy of the mean isoform expression proportions across all cells, normalised by the maximum possible entropy as

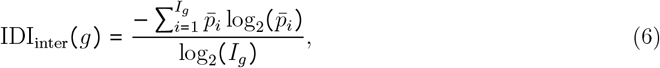

where 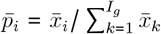, and 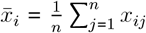 is the mean expression of isoform *i* across all cells. This metric quantifies the overall diversity of isoforms used across the entire cell population.

#### III: Intra-cell-type heterogeneity

To quantify cell-to-cell variation in isoform usage within a given cell type, `ScIsoX` computes the average Jensen-Shannon distance between cells. For a gene *g* ∈ *𝒢* expressed in cell type *𝒮*_*k*_ ∈ *𝒮* with *n*_*k*_ cells, intra-cell-type heterogeneity is defined as

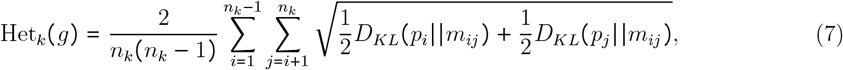

where *p*_*i*_ and *p*_*j*_ are the isoform proportion vectors for cells *c*_*i*_ ∈ 𝒞 and *c*_*j*_ ∈ 𝒞 in the cell type 𝒮_*k*_, *m*_*ij*_ is their average distribution, and *D*_*KL*_ is the Kullback-Leibler divergence. The overall intra-cell-type heterogeneity is calculated as the mean across all cell types where the gene is expressed. This metric measures cell-to-cell variation in isoform usage within each cell type. It reveals whether cells of the same type use isoforms consistently.

#### IV: Inter-cell-type specificity

To assess how distinctly a gene deploys its isoforms across different cell types, `ScIsoX` computes the average Jensen-Shannon distance between cell-type-specific isoform profiles. For a gene *g* ∈ 𝒢 expressed in *S* cell types, inter-cell-type specificity is defined as

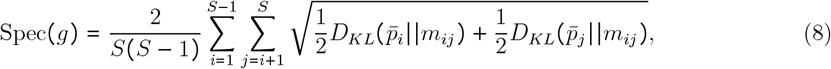

where 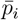 and 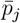 are the vectors of mean isoform proportions for cell types 𝒮_*i*_ ∈ 𝒮 and 𝒮_*j*_ ∈ 𝒮. Higher values indicate cell-type-specific isoform usage patterns, suggesting specialised functional roles across different cell populations.

#### V: Intra-cell-type heterogeneity variability

To determine whether cellular heterogeneity is concentrated in specific cell types, `ScIsoX` computes the coefficient of variation (CV) of intra-cell-type heterogeneity values across cell types. For a gene *g* ∈ *𝒢* expressed in *S* cell types, this variability is defined as

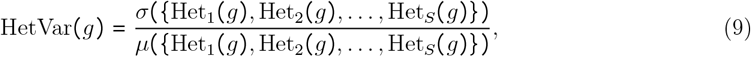

where *σ* and *μ* represent the standard deviation and mean, respectively. High values indicate that some cell types have higher internal heterogeneity than others, suggesting targeted subpopulation structure or regulatory plasticity within specific lineages.

#### VI: Inter-cell-type difference variability

To assess whether isoform usage differences are concentrated between specific cell type pairs, `ScIsoX` computes the CV of pairwise Jensen-Shannon distances. For a gene *g* ∈ *𝒢* expressed in *S* cell types, this variability is defined as

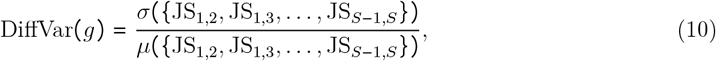

where JS_*i,j*_ is the Jensen-Shannon distance between cell types *i* and *j*. High values indicate that certain cell type pairs exhibit particularly divergent isoform usage patterns, suggesting lineage-specific splicing regulation or functional specialisation between specific cell populations.

#### VII: Cell-type-specific co-expression variability

To evaluate whether a gene is subject to different co-expression patterns across cell types, `ScIsoX` computes the CV of mean intra-cellular diversity across cell types. For a gene *g* ∈ 𝒢 expressed in *S* cell types, this variability is defined as

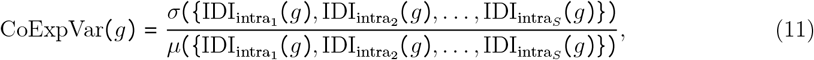

where 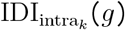 is the mean intra-cellular isoform diversity of gene *g* in cell type 𝒮_*k*_ (i.e., the tendency for genes to co-express multiple isoforms across cell type 𝒮_*k*_). High values indicate that a gene exhibits dramatically different co-expression patterns in different cellular contexts, suggesting context-dependent regulation of isoform co-expression.

#### Additional complexity metrics

`ScIsoX` computes a range of supplementary metrics to further characterise isoform expression pattern. These additional metrics are detailed in (Supplementary Table 2).

### Optimal threshold determination for complexity classification

The `ScIsoX` framework implements an advanced multi-stage statistical pipeline to determine optimal classification thresholds for each complexity dimension. This methodology addresses the challenges of analysing heterogeneous distribution patterns observed in transcriptomic complexity metrics.

#### Distribution-aware preprocessing

For each complexity metric, we first apply distribution-aware preprocessing to identify the underlying distribution characteristics. This preprocessing phase employs multiple statistical approaches:

1. **Distribution classification:** Each metric’s distribution is classified into one of several categories: multimodal, zero-inflated, extremely skewed, moderately skewed, or unimodal using a comprehensive multi-method approach. For multimodality detection, we employ three complementary methods, namely Hartigan’s dip test for statistical significance, kernel density estimation with adaptive bandwidth selection for peak/valley analysis, and Gaussian mixture modelling with Bayesian Information Criterion for component separation. Skewness is assessed using moment-based calculations with distinct thresholds for moderate and extreme cases.
2. **Zero-inflation detection:** An adaptive histogram-based approach is used to identify zero-inflated distributions. This method calculates a data-dependent near-zero threshold (based on range and interquartile range), determines optimal bin width using the Freedman-Diaconis rule, and analyses the ratio between first and second bins to detect significant zero-inflation. For identified zero-inflated distributions, we further characterise the non-zero component, testing for multimodality and skewness to determine appropriate transformation strategies.
3. **Transformation application:** When necessary, Yeo-Johnson transformations are applied with optimised parameters to normalise extremely skewed distributions while preserving their essential characteristics for threshold determination.

#### Distribution-specific threshold algorithms

Based on the identified distribution type, specialised algorithms are employed to determine optimal thresholds, with a hierarchical fallback strategy to ensure robust results (Supplementary Fig.7-8):

1. **For multimodal distributions:** our algorithm first attempts to identify inflection points in the density curve, followed by mixture model-based component separation if needed. For distributions with clear valleys between modes, it calculates the optimal separation threshold based on relative depths and positions of these valleys.
2. **For extremely skewed distributions:** our algorithm avoids extreme tails by focusing on the central mass of the distribution, using inflection point and curvature analysis to identify natural separation points.
3. **For zero-inflated distributions:** the non-zero component is extracted and analysed separately, using either gap detection (for significant discontinuities), mixture modelling (for multimodal non-zero components), or adaptive percentiles based on the skewness of the non-zero component.
4. **For moderately skewed and unimodal distributions:** our algorithm employs a combination of density curve analysis, distribution moments, and weighted mixtures of normal distributions to identify optimal decision boundaries.

Each method includes reliability assessment, with automatic fallback to simpler techniques when necessary. This adaptive approach ensures robust threshold determination across diverse distribution patterns encountered in complexity metrics.

#### Statistical validation framework

The reliability of determined thresholds is assessed through a comprehensive validation framework:

1. **Bootstrap stability assessment:** `ScIsoX` performs 100 (adjustable) bootstrap iterations, recalculating the threshold for each resampled dataset. This provides confidence intervals, standard deviations, and coefficients of variation that inform reliability scores, with higher weight given to stable thresholds.
2. **K-fold cross-validation:** For datasets with sufficient samples, `ScIsoX` performs stratified k-fold cross-validation to assess threshold consistency across different subsets of the data. The cross-validation coefficient of variation is integrated into the final reliability assessment.
3. **Distribution-specific reliability adjustment:** Initial reliability scores derived from the primary threshold method are adjusted based on distribution characteristics, with higher penalties for problematic distributions and distributions with limited supporting data.
4. **Sanity checking:** Final thresholds undergo verification against the data distribution’s quantiles to ensure they are reasonable, with automated adjustments applied when necessary to prevent threshold placement in extreme distribution tails.

### Classification system

Based on the seven core metrics, we developed a multi-dimensional classification system that categorises genes according to their complexity profiles (Supplementary Table 3). For each dimension, genes are classified into biologically meaningful categories based on the thresholds derived from our distribution-specific threshold algorithms:

1. Intra-cellular isoform diversity is classified as “Strong Isoform Co-expression” or “Weak Isoform Co-expression”
2. Inter-cellular isoform diversity is classified as “High Isoform Diversity” or “Low Isoform Diversity” (see for example in Fig.1b)
3. Intra-cell-type heterogeneity is classified as “High Cellular Heterogeneity” or “Low Cellular Heterogeneity”
4. Inter-cell-type specificity is classified as “Cell-Type-Specific Isoform Expression” or “Cell-Type-Independent Isoform Expression” (see for example in Fig.1b)
5. Intra-cell-type heterogeneity variability is classified as “Variable Heterogeneity Across Cell Types” or “Consistent Heterogeneity Across Cell Types”
6. Inter-cell-type difference variability is classified as “High Cell-Type Distinctions” or “Low Cell-Type Distinctions”
7. Cell-type-specific co-expression variability is classified as “Cell-Type-Adaptive Co-expression” or “Cell-Type-Consistent Co-expression”

NA values are preserved throughout this process, as they represent biologically meaningful cases such as single-isoform genes for diversity metrics or genes expressed in only one cell type for inter-cell-type metrics. The integrated classification system enables systematic comparison of transcriptomic complexity patterns across genes and facilitates the identification of genes with interesting or unusual complexity profiles (see example in Supplementary Table 4).

### Visualisation and Analysis Features

The `ScIsoX` framework implements a comprehensive suite of visualisation and analysis tools designed to explore and interpret multidimensional transcriptomic complexity patterns. The framework’s data structure facilitates efficient analytical workflows that enable researchers to gain biological insights from complex isoform expression patterns.

#### Core Data Structures and Organisation

The framework organises isoform complexity data into two complementary object structures that support diverse analytical approaches. The `IntegratedSCHT` object encapsulates gene-level isoform expression matrices in a hierarchical structure, with both global and cell-type-specific expression patterns stored efficiently in a list-based format. The `transcriptomic_complexity` object contains a data frame of complexity metrics (`metrics`), cell-type-specific measurements (`cell_type_metrics`), classification thresholds, and statistical metadata. These data structures, combined with the S3 method system for object manipulation in `R`, enable sophisticated data exploration.

#### Analytical Tools

1. **Cell-Type-Specific Complexity Analysis** calculates and compares complexity metrics independently for each cell type, enabling the identification of cell types with distinctive isoform regulation patterns.
2. **Complexity Pattern Filtering** identifies genes matching specific combinations of complexity classifications across multiple dimensions using the `find_complexity_pattern()` function. This enables targeted discovery of genes with precise complexity signatures of interest.
3. **Gene Selection Tool** extracts genes with specific complexity characteristics using the `select_genes_of_interest()` function with customisable filtering criteria.
4. **Complexity Metric Comparison** extracts and compares transcriptomic complexity metrics across multiple genes using the `compare_gene_metrics()` function for custom visualisations or statistical analyses.

#### Visualisation capabilities

1. **Complexity Landscape Visualisations** generate bivariate scatter plots that position genes across two complexity dimensions with integrated marginal distributions. These plots incorporate quadrant statistics and threshold lines to identify genes with exceptional complexity profiles (Fig.1b Top). Interactive highlighting capabilities facilitate the identification of notable genes.
2. **Density Contour Maps** overlay kernel density estimation contours on complexity landscapes to reveal clustering patterns and high-density regions in the complexity space (Fig.1b Bottom). Density contour maps employ adaptive bandwidth algorithms that accommodate varying data densities and highlight regions of biological significance through smooth visualisation of gene concertration hotspots.
3. **Complexity Radar Charts** visualise the complete seven-dimensional complexity signature of individual genes or comparative profiles across multiple genes (Fig.2e and Supplementray Fig.3a). The implementation supports various normalisation methods and custom axis configurations for effective comparison of complexity profiles.
4. **Multi-gene Cell-Type-specific Radar Charts** facilitate the comparison of complexity profiles across multiple genes and cell types in a structured grid layout, enabling the identification of cell-type-specific regulatory patterns (Supplementray Fig.3b). The implementation includes options for global or per-cell type scaling.
5. **Dual Diversity Plots** are scatter plots for intra-cellular and inter-cellular diversity metrics with diagonal reference lines indicating the theoretical equality boundary (Fig.2a). This visualisation specifically highlights genes exhibiting unusual diversity patterns, which may indicate specialised regulatory mechanisms.
6. **Co-expression Correlation Heatmaps** are hierarchically clustered correlation matrices of isoform expression patterns, automatically detecting modules of coordinated or mutually exclusive isoform usage (Fig.2b). The implementation allows users to select among multiple correlation methods (Pearson, Spearman, and Kendall) to accommodate different data distributions and analytical requirements, with options to display correlation values directly. These heatmaps are generated using the ComplexHeatmap package [11].
7. **Isoform Usage Profile Plots** are stacked bar charts that display proportions of expressed isoform usage across cell types, developmental stages, or experimental conditions (Fig.2c). These plots include automatic minor isoform grouping and customisable cell type ordering, facilitating the identification of cell-type-specific isoform preferences.
8. **Isoform Transition Plots** visualise dynamic changes in isoform usage across ordered cell types, time points, or developmental stages (Fig.2d). This approach is particularly effective for revealing isoform switching events during differentiation processes or disease progression.
9. **Ridge Plots** visualise the distribution of complexity metrics through overlapping density curves (Supplementary Fig.3). The implementation supports both global complexity comparisons across metrics and cell-type-specific analyses, offering a compact way to compare multiple distributions simultaneously.
10. **Distribution Threshold Fitting Plots** visualise the distributions of complexity metrics across multiple cell types with optimal threshold determined by the algorithm (Supplementary Fig.7 - 8).
11. **Complexity Metric Heatmaps** provide a comprehensive view of multiple complexity metrics across different groups or conditions (Fig.2f). These heatmaps can be configured to show absolute values or changes between consecutive conditions, with gene selection based on variance, magnitude of change, or custom gene lists. These heatmaps are generated using the ComplexHeatmap package [11].
12. **Group Comparison Density Difference Maps** calculate and visualise the density differences of genes in 2D metric space between different experimental groups or conditions (Supplementary Fig.5 - 6). These visualisations help identify regions where gene distributions shift across conditions, revealing patterns of coordinated complexity changes in response to experimental manipulations.

All analysis and visualisation functions support comprehensive parameterisation while maintaining computational efficiency for large-scale datasets.

### Cell type annotation

All datasets used in this study include cell type annotations acquired through different methodologies. For the murine haematopoietic development dataset, cell type annotations were based on experimentally validated labels [4]. For the brain dataset, we utilised preprocessed data and annotations from the study [5]. In that study, computational preprocessing was performed using Seurat [12], and annotations were generated through manual marker gene identification. This approach identified major brain cell populations including excitatory and inhibitory neurons, oligodendrocytes, astrocytes, microglia, vascular cells, and progenitor populations. For the PBMC dataset, we also performed preprocessing using Seurat [12] and subsequently classified cell types computationally using SingleR [13]. The reference dataset contained well-characterised immune cell signatures that enabled identification of T cells, B cells, NK cells, monocytes, and other PBMC subpopulations.

### Considerations and limitations

The structured organisation of complexity metrics and hierarchical tensor format facilitates integration with complementary single-cell analysis approaches. The quantitative metrics can be correlated with differential expression patterns to identify relationships between expression levels and isoform regulation mechanisms, allowing researchers to relate changes in transcriptomic complexity with expression level alterations across conditions. The transcriptomic complexity signatures can also be correlated with DNA binding motif enrichment patterns to identify potential regulatory elements driving specific complexity profiles. Moreover, the framework’s cell type-resolved metrics can be mapped onto trajectory inference results, e.g., to characterise dynamic changes in isoform usage mechanisms during cellular differentiation processes. The classification system enables the incorporation of complexity dimensions into gene regulatory network analyses, potentially revealing how splicing regulators influence network topology and dynamics. Furthermore, these metrics support cross-species comparisons to investigate evolutionary conservation of isoform regulation patterns.

Several limitations should be considered when applying `ScIsoX`. First, some metrics depend on accurate cell type annotation. While the framework is compatible with any popular single-cell clustering and annotation method, the quality of cell type definitions will affect the accuracy of specific metrics, particularly those based on cell type comparisons. In cases where cell type boundaries are ambiguous or annotations uncertain, users should exercise caution when interpreting results or focus on metrics that do not depend on cell type information.

Additionally, users should note that the final number of HVGs in the created SCHT structure may differ from the initial *n*_hvg_ parameter specified in the `create_scht()` function. This discrepancy occurs because the algorithm removes genes with only a single isoform after processing, ensuring that all genes included in subsequent analyses have multiple expressed isoforms. This filtering step is essential for meaningful isoform complexity analysis but may reduce the final gene count. Improved sequencing quality and depth can significantly mitigate this issue by enabling more comprehensive isoform detection. If researchers wish to maximise the number of genes in downstream analyses, they can increase the *n*_hvg_ parameter, though this value cannot exceed the total number of genes present in the dataset.

While the hierarchical data structure offers computational advantages for typical single-cell datasets, extremely large datasets may still require additional optimisation strategies. The framework includes options for batch-wise processing and memory-efficient data handling to address these scenarios.

## Data Availability

No datasets were generated during the current study.

## Code Vailability

`ScIsoX` is available under the MIT License at https://github.com/ThaddeusWu/ScIsoX

## Acknowledgements

U.S. received support from the National Health and Medical Research Council (Investigator Grant #1196405), the Tropical Australian Academic Health Centre (Project Grant SF01124), and the Townsville University Hospital (Grant #THHSSERTA_RPG05_2024, #THHSSERTA_RPG15_2024, and #THHSSERTA_RCG05_2024).

## Author Contributions

S.W. and U.S. conceived the topic and structure. S.W. developed the package and drafted the manuscript. U.S. supervised the work and drafted the manuscript. Both authors revised the paper and approved the final version.

