## Supplementary Information for "ScIsoX: A Multidimensional Framework for Measuring Isoform-Level Transcriptomic Complexity in Single Cells"

Siyuan Wu[1,2,3] and Ulf Schmitz[1,2,*]

[1]Computational Biomedicine Lab, College of Science and Engineering, James Cook University, Townsville, Queensland, Australia

[2]Centre for Tropical Bioinformatics and Molecular Biology, James Cook University, Cairns, Queensland, Australia

[3]School of Mathematics, Monash University, Melbourne, Victoria, Australia

May 11, 2025

| Metric | Description | Calculation Method | Biological Interpretation | High Values Indicate | Low Values Indicate |
|---|---|---|---|---|---|
| **Intra-cellular Isoform Diversity** | Measures the tendency for a gene to co-express multiple isoforms within individual cells | Weighted average of per-cell normalised Shannon entropy | Reveals whether genes express multiple isoforms simultaneously | Active co-expression of multiple isoforms within individual cells (potential functional cooperation) | Expression of predominantly one isoform per cell (potential mutually exclusive regulation) |
| **Inter-cellular Isoform Diversity** | Quantifies the diversity of isoforms expressed by a gene across the whole cell population | Normalised Shannon entropy of average isoform expression | Shows the overall diversity of isoforms expressed in the tissue | Multiple isoforms present across the cell population (diverse functionality) | One or few dominant isoforms across the population (consistent functionality) |
| **Intra-cell-type Heterogeneity** | Measures cell-to-cell variation in isoform usage within each cell type | Average Jensen-Shannon distance between cells within a cell type | Reveals whether cells of the same type use isoforms consistently | High cell-to-cell variability in isoform usage (potential sub-populations or states) | Consistent isoform usage pattern within the cell type (consistent functionality) |
| **Inter-cell-type Specificity** | Quantifies how differently a gene uses its isoforms across different cell types | Average Jensen-Shannon distance between cell type-specific isoform profiles | Shows whether isoform usage is cell-type-specific | Cell types use distinct isoform repertoires (specialised functions) | Similar isoform usage across cell types (conserved functions) |
| **Intra-cell-type Heterogeneity Variability** | Measures whether certain cell types show particularly high cellular heterogeneity | Coefficient of variation of heterogeneity values across cell types | Reveals if heterogeneity is concentrated in specific cell types | Some cell types have much higher internal heterogeneity than others (targeted sub-population structure) | Consistent levels of heterogeneity across all cell types (uniform regulation) |
| **Inter-cell-type Difference Variability** | Measures whether certain cell type pairs show particularly significant differences | Coefficient of variation of pairwise Jensen-Shannon distances | Shows whether differences are concentrated between specific cell type pairs | Certain cell type pairs have dramatically different isoform usage (lineage-specific divergence) | Relatively uniform differences between all cell type pairs (gradual differentiation) |
| **Cell-type-specific Co-expression Variability** | Measures whether a gene exhibits different co-expression patterns in different cell types | Coefficient of variation of mean intra-cellular diversity across cell types | Reveals if co-expression patterns vary by context | Gene exhibits different co-expression patterns in different cell types (context-dependent regulation) | Consistent co-expression mechanism across all cell types (fixed regulatory mechanism) |

Supplementary Table 1: **Core Transcriptomic Complexity Metrics**

| Metric | Description | Calculation Method | Biological Interpretation |
|---|---|---|---|
| **IDI Difference** | The difference between inter-cellular and intra-cellular isoform diversity | Inter-cellular isoform diversity – Intra-cellular isoform diversity | Positive values indicate greater diversity across cells than within cells, suggesting cell specialisation; negative values suggest cells co-express many isoforms but in similar patterns |
| **Simpson Index** | Alternative diversity measure more sensitive to dominant isoforms | 1 - sum(iso_props$^2$) | Accounts for the probability that two randomly selected transcripts belong to different isoforms; complements Shannon entropy-based metrics |
| **Evenness** | Normalised diversity controlling for number of expressed isoforms | inter_cellular_isoform_diversity / log2(n_expressed_isoforms) | Measures how equally expressed the isoforms are; values near 1 indicate equal expression, near 0 indicate dominance by few isoforms |
| **Dominant Isoform Proportion** | The proportion of the most abundant isoform | max(iso_props) | Indicates the degree of dominance by a single isoform; high values suggest one functional isoform dominates |
| **Number of Expressed Isoforms** | Count of isoforms with detectable expression | sum(iso_means > 0) | Indicates the absolute diversity of isoforms being expressed; reflects the gene's splicing complexity |
| **Percentage of Multi-isoform Cells** | Percentage of cells expressing multiple isoforms of a given gene | (Multi-isoform cell count / cells expressing) × 100 | Direct measure of co-expression at the single-cell level; high values indicate widespread co-expression |
| **Cells Expressing** | Number of cells with detectable expression of a given gene | sum(cell_sums > 0) | Indicates the prevalence of the gene's expression across the cell population |
| **Percentage of Cells Expressing** | Percentage of total cells expressing the gene | (cells_expressing / total_cells) × 100 | Measures how widespread the gene's expression is; distinguishes ubiquitous from restricted expression patterns |

Supplementary Table 2: **Additional Transcriptomic Complexity Metrics**

| Dimension | High Classification | Low Classification | Interpretation |
|---|---|---|---|
| **Intra-cellular Isoform Diversity** | High Isoform Co-expression | Low Isoform Co-expression | Distinguishes genes based on single-cell co-expression patterns |
| **Inter-cellular Isoform Diversity** | High Isoform Diversity | Low Isoform Diversity | Distinguishes genes based on population-level isoform diversity |
| **Intra-cell-type Heterogeneity** | High Cellular Heterogeneity | Low Cellular Heterogeneity | Distinguishes genes based on cell-to-cell variability within cell types |
| **Inter-cell-type Specificity** | Cell-Type-Specific Isoform Expression | Cell-Type-Independent Isoform Expression | Distinguishes genes based on cell type specialisation of isoform usage |
| **Intra-cell-type Heterogeneity Variability** | Variable Heterogeneity Across Cell Types | Consistent Heterogeneity Across Cell Types | Distinguishes genes based on targeted vs. uniform heterogeneity |
| **Inter-cell-type Difference Variability** | High Cell-Type Distinctions | Low Cell-Type Distinctions | Distinguishes genes based on focused vs. gradual differentiation |
| **Cell-type-specific Co-expression Variability** | Cell-Type-Adaptive Co-expression | Cell-Type-Consistent Co-expression | Distinguishes genes based on context-dependent isoform co-expression |

Supplementary Table 3: **Metrics Classification System**

| Pattern | Biological Significance | Example Genes | Potential Functional Implications |
|---|---|---|---|
| **High Intra-cellular Isoform Diversity + High Inter-cellular Isoform Diversity** | Rich isoform landscape both within and across cells | Genes involved in complex cellular processes | Multiple functional isoforms with complementary roles; high regulatory complexity |
| **Low Intra-cellular Isoform Diversity + High Inter-cellular Isoform Diversity** | Cell specialisation in isoform usage | Cell type marker genes, specialised receptors | Cell type-specific isoform selection; potential for specialised functions |
| **Low Intra-cellular Isoform Diversity + Low Inter-cellular Isoform Diversity** | Single dominant isoform usage | Housekeeping genes, core cellular machinery | Conserved function requiring specific isoform; limited need for diversity |
| **High Intra-cellular Isoform Diversity + Low Inter-cellular Isoform Diversity** | Consistent co-expression of specific isoform sets | Genes requiring balanced isoform ratios | Functional requirement for multiple isoforms within same cell; potential isoform cooperation |
| **High Cell-type Specificity + Low Difference Variability** | Consistent differentiation across all cell types | Lineage-specific transcription factors | Gradual divergence in isoform usage corresponding to cellular differentiation |
| **High Cell-type Specificity + High Difference Variability** | Targeted differentiation between specific lineages | Immune recognition molecules, neuronal connectivity factors | Sharp transitions in isoform usage between specific lineages; potential for dramatic functional shifts |

Supplementary Table 4: **Example of Complexity Patterns**

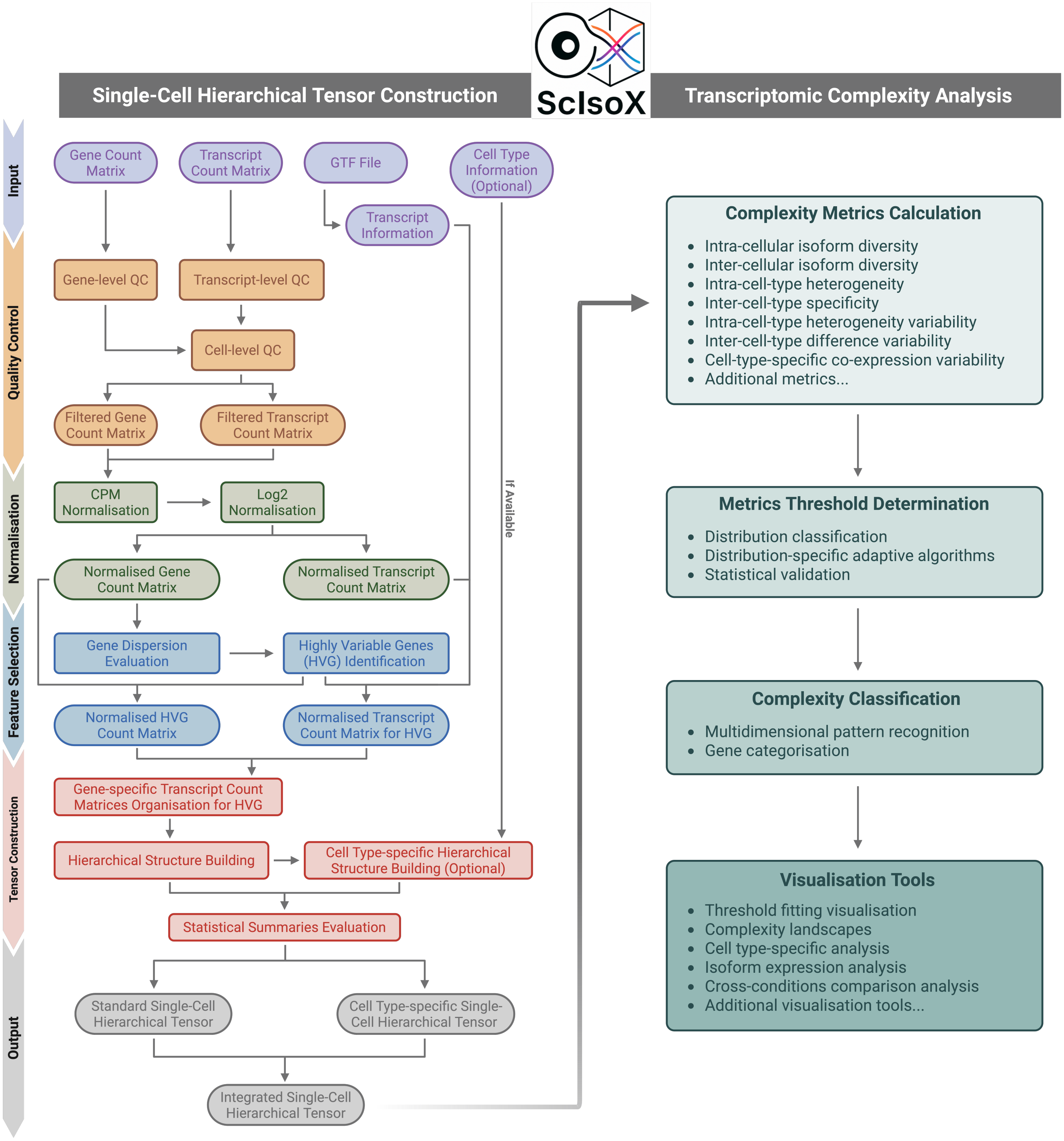


Supplementary Figure 1: **Detailed workflow of the ScIsoX analysis framework.** This is the comprehensive overview of the ScIsoX analysis pipeline, beginning with input data on the left (gene count matrix, transcript count matrix, GTF file, and cell type information), proceeding through quality control, normalisation, highly variable genes selection, SCHT construction, and culminating in the various components of transcriptomic complexity analysis on the right. The transcriptomic complexity analysis includes complexity metrics calculation (seven core metrics and additional metrics), distribution-based adaptive threshold determination, multidimensional complexity classification, and rich visualisation tools. Each processing step is clearly defined, forming a complete analytical pipeline that supports extraction of biological meaning from SCLRS data. Created with BioRender.com.

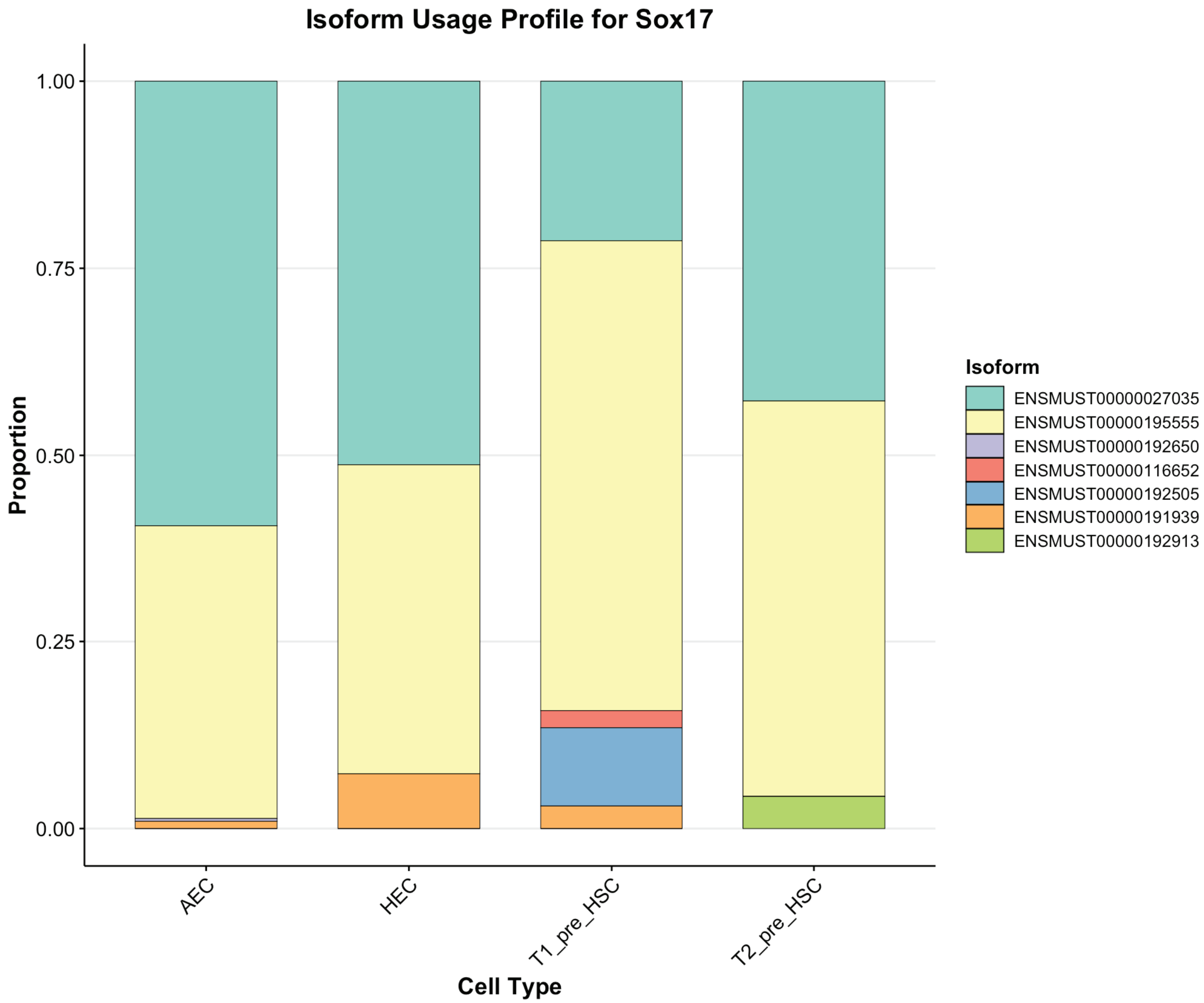


Supplementary Figure 2: ***Sox17* isoform usage profiles across different mouse haematopoietic developmental stage.** AEC - arterial endothelial cells; HEC - hemogenic endothelial cells; T1__pre__HSC - Type 1 precursor hematopoietic stem cells; T2__pre__HSC - Type 2 precursor hematopoietic stem cells.

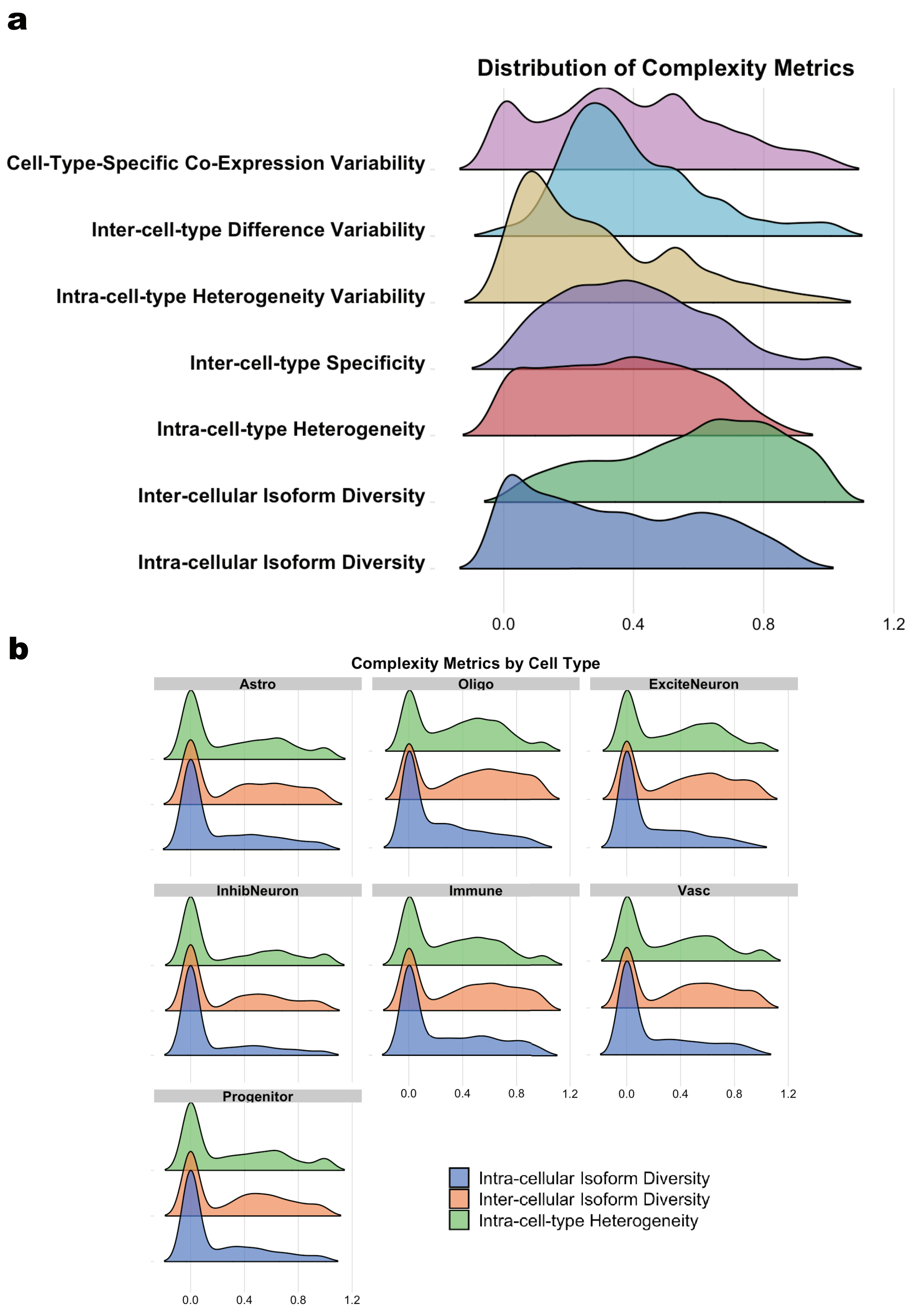


Supplementary Figure 3: **Distribution characteristics of the seven core complexity metrics and cell-type-specific patterns.** (a) Ridge plots showing distributions of the seven core complexity metrics in the brain dataset, displaying characteristic distributions for each dimension. Each metric exhibits a unique distribution profile, reflecting the biological complexity captured by different dimensions. (b) Comparison of density plots for three key complexity metrics across seven cell types in the brain dataset. Immune - immune cells; Astro - astrocytes; Oligo - oligodendrocytes; ExciteNeuron - excitatory neurons; InhibNeuron - inhibitory neurons; Progenitor - progenitor cells; Vasc - vascular cells.

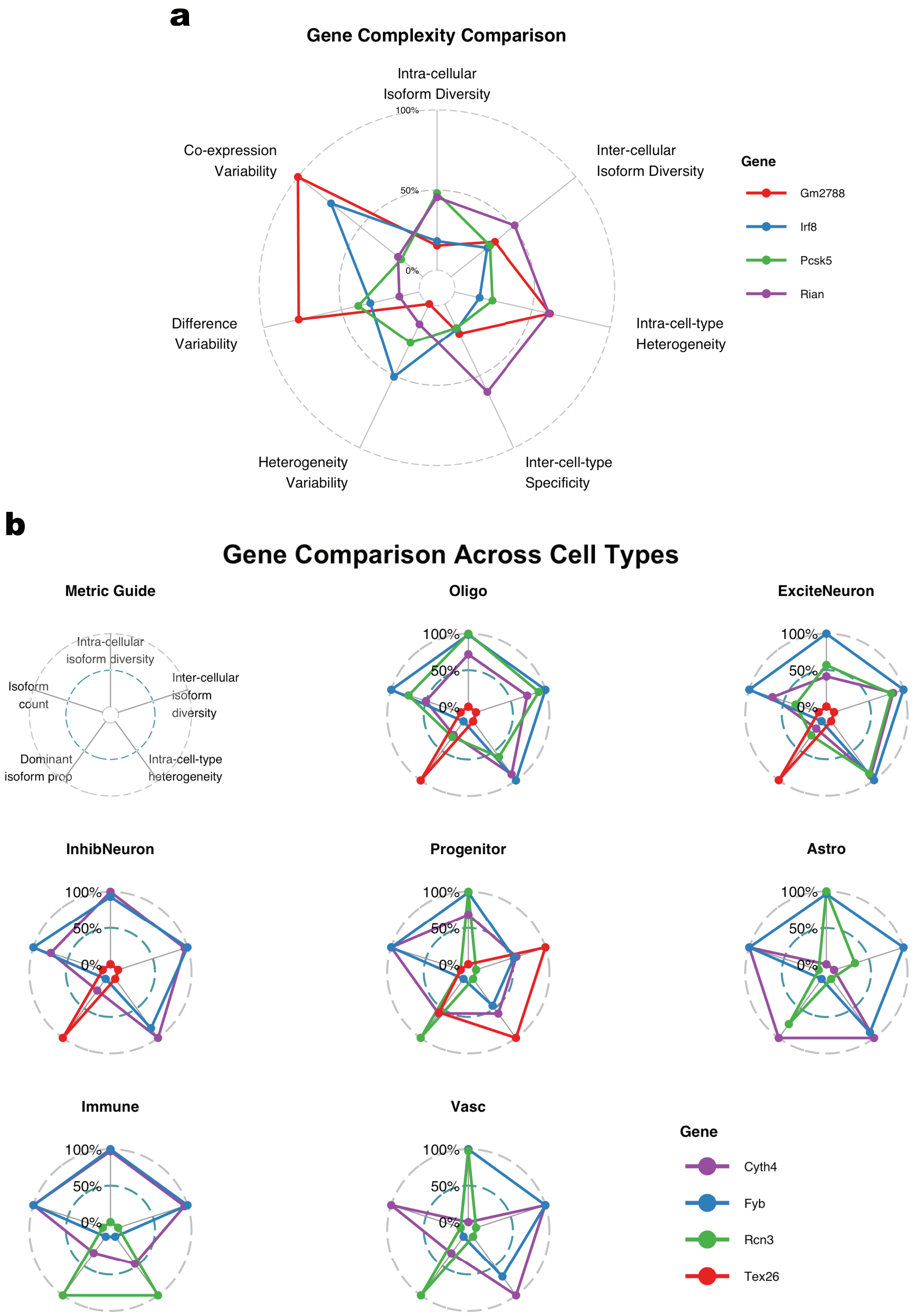
a
Gene Complexity Comparison
Intra-cellular
Isoform Diversity
Inter-cellular
Isoform Diversity
Intra-cell-type
Heterogeneity
Inter-cell-type
Specificity
Heterogeneity
Variability
Difference
Variability
Co-expression
Variability
100%
50%
0%
Gene
Gm2788
Irf8
Pcsk5
Rian
b
Gene Comparison Across Cell Types
Metric Guide
Intra-cellular
isoform diversity
Inter-cellular
isoform
diversity
Intra-cell-type
heterogeneity
Dominant
isoform prop
Isoform
count
Oligo
ExciteNeuron
InhibNeuron
Progenitor
Astro
Immune
Vasc
100%
50%
0%
Gene
Cyth4
Fyb
Rcn3
Tex26

Supplementary Figure 4: **Radar chart visualisations for multidimensional complexity comparisons.** (a) Seven-dimensional complexity metric comparison radar charts for four genes (*Gm2788*, *Irf8*, *Pcat5*, and *Ran*) in the mouse early blood development dataset, showcasing the unique complexity signatures of different genes. Each gene displays a distinctive complexity fingerprint, reflecting its specific isoform regulatory mechanisms. (b) Complexity profiles of individual genes (*Fyb*, *Mxra8*, *Rcn3*, and *Tex26*) across eight different brain cell types visualised with radar charts, demonstrating cell type specificity of isoform regulation. A metric guide in the top-left corner provides the meaning of the five axes in the radar charts. This visualisation method effectively captures and compares high-dimensional complexity data. Immune - immune cells; Astro - astrocytes; Oligo - oligodendrocytes; ExciteNeuron - excitatory neurons; InhibNeuron - inhibitory neurons; Progenitor - progenitor cells; Vasc - vascular cells.

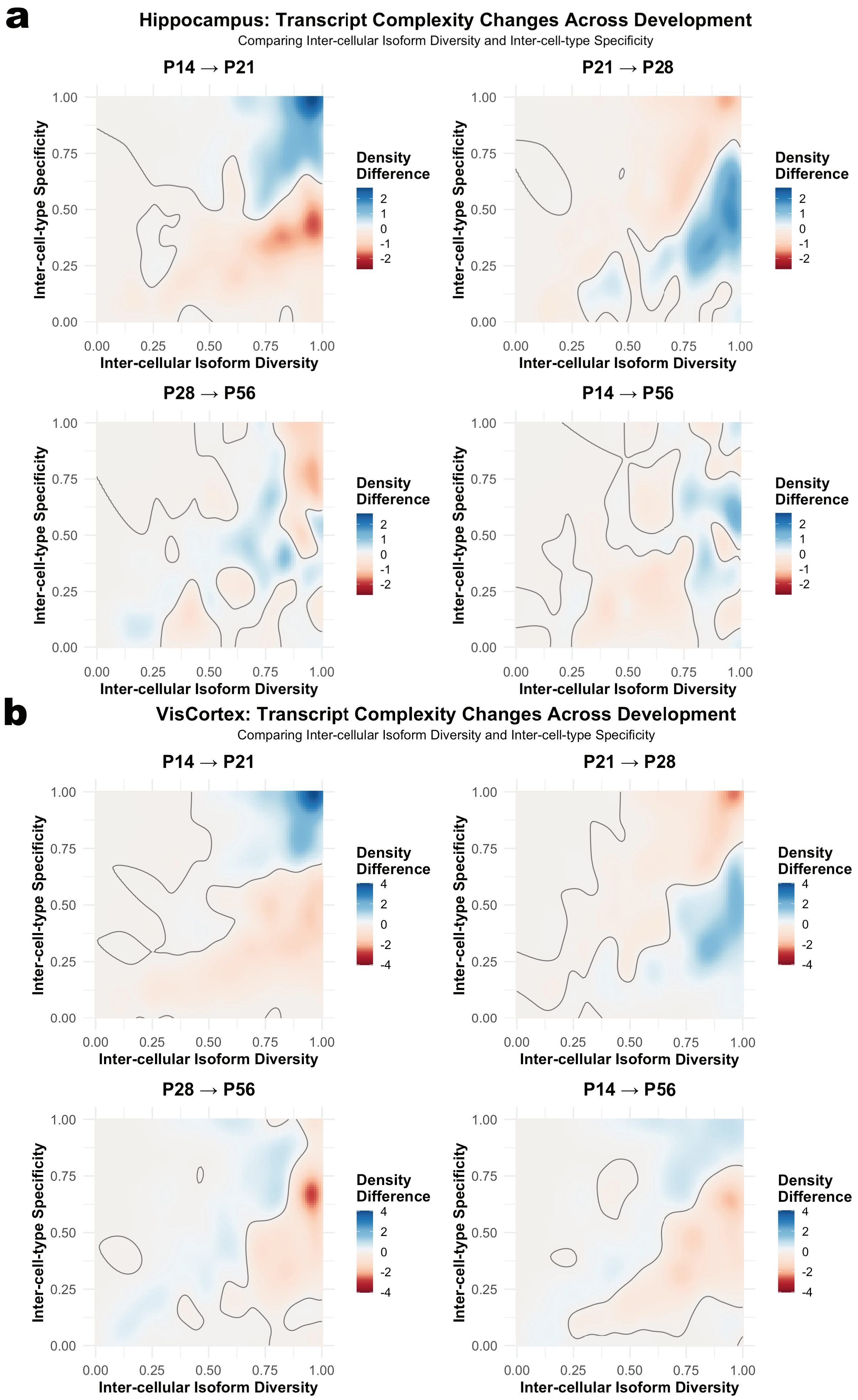
a
Hippocampus: Transcript Complexity Changes Across Development
Comparing Inter-cellular Isoform Diversity and Inter-cell-type Specificity
P14 → P21
P21 → P28
P28 → P56
P14 → P56
Inter-cell-type Specificity
Inter-cellular Isoform Diversity
0.00
0.25
0.50
0.75
1.00
Density Difference
2
1
0
-1
-2
b
VisCortex: Transcript Complexity Changes Across Development
Comparing Inter-cellular Isoform Diversity and Inter-cell-type Specificity
P14 → P21
P21 → P28
P28 → P56
P14 → P56
Inter-cell-type Specificity
Inter-cellular Isoform Diversity
Density Difference
4
2
0
-2
-4

Supplementary Figure 5: **Dynamic changes in transcriptomic complexity across postnatal developmental stages in mouse hippocampus and visual cortex regions.** (a) Density difference maps comparing inter-cellular isoform diversity and inter-cell-type specificity in mouse hippocampus across four developmental stages (Days 14, 21, 28 and 56). Red regions indicate gene decreasing density while blue regions indicate increasing density. Each transition period shows a unique pattern of changes. (b) Complexity change patterns in mouse visual cortex across four developmental stages, showing region-specific differences compared to hippocampus. Colour scales indicate the intensity of density differences. This analysis reveals spatiotemporal changes in transcriptomic complexity during brain development.

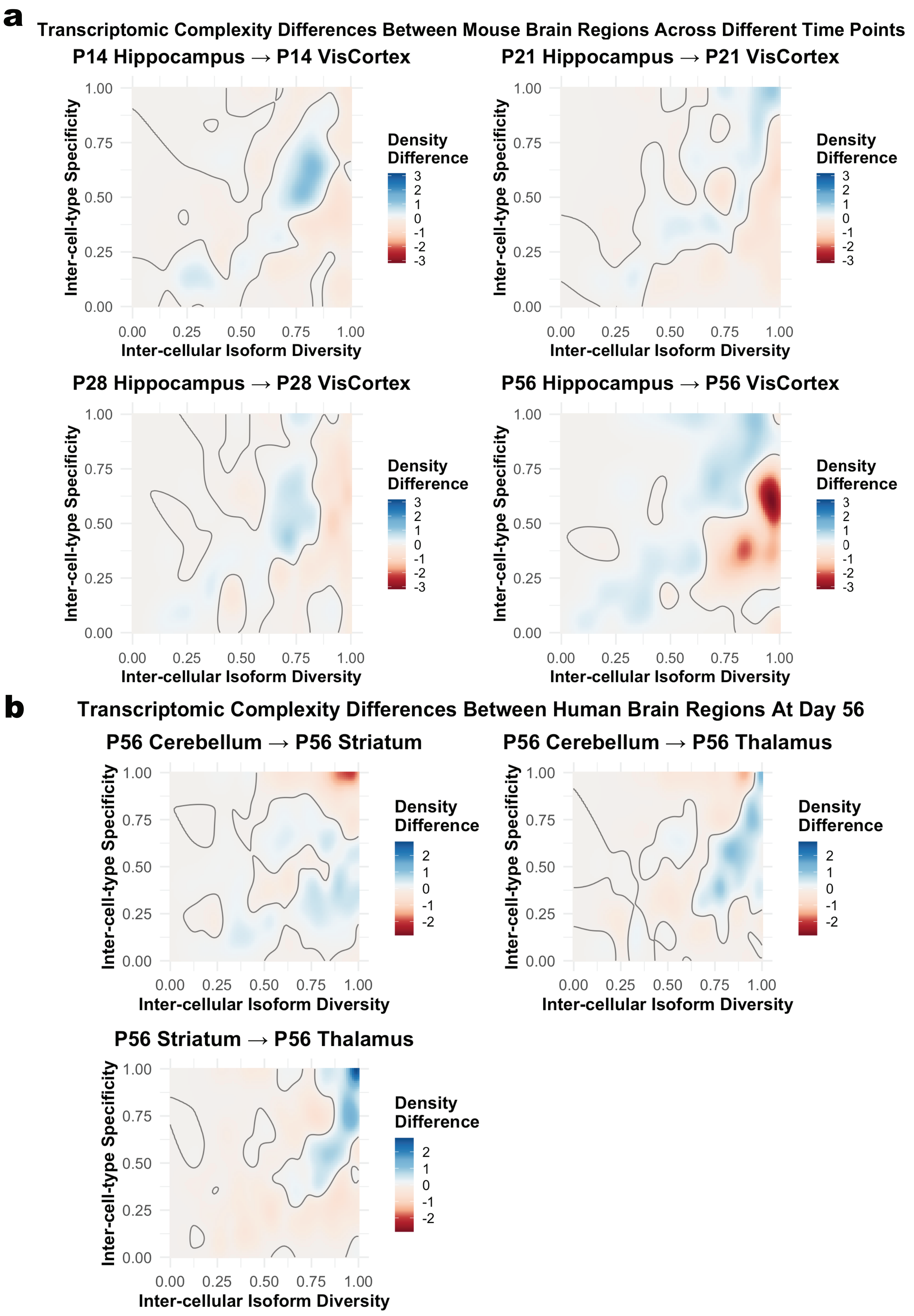
a
Transcriptomic Complexity Differences Between Mouse Brain Regions Across Different Time Points
P14 Hippocampus → P14 VisCortex
P21 Hippocampus → P21 VisCortex
P28 Hippocampus → P28 VisCortex
P56 Hippocampus → P56 VisCortex
Inter-cell-type Specificity
Inter-cellular Isoform Diversity
0.00
0.25
0.50
0.75
1.00
Density Difference
3
2
1
0
-1
-2
-3
b
Transcriptomic Complexity Differences Between Human Brain Regions At Day 56
P56 Cerebellum → P56 Striatum
P56 Cerebellum → P56 Thalamus
P56 Striatum → P56 Thalamus
Inter-cell-type Specificity
Inter-cellular Isoform Diversity
0.00
0.25
0.50
0.75
1.00
Density Difference
2
1
0
-1
-2

Supplementary Figure 6: **Transcript complexity differences between mouse brain regions across postnatal developmental stages and between adult human brain regions.** (a) Developmental changes in transcript complexity profiles comparing hippocampus and visual cortex regions across different time points in mouse brain (Days 14, 21, 28 and 56). Each density difference map displays changes in the relationship between inter-cellular isoform diversity and inter-cell-type specificity, with blue indicating regions of increased gene density and red indicating decreased density. These developmental comparisons reveal dynamic shifts in transcriptomic complexity patterns as brain circuits mature. (b) Transcript complexity differences between adult human brain regions. The figure presents density difference maps between three pairs of distinct brain regions (Cerebellum-Striatum, Cerebellum-Thalamus, and Striatum-Thalamus). Each comparison displays regional differences in inter-cellular isoform diversity and inter-cell-type specificity, with blue indicating greater gene density in the first region and red indicating greater density in the second region.

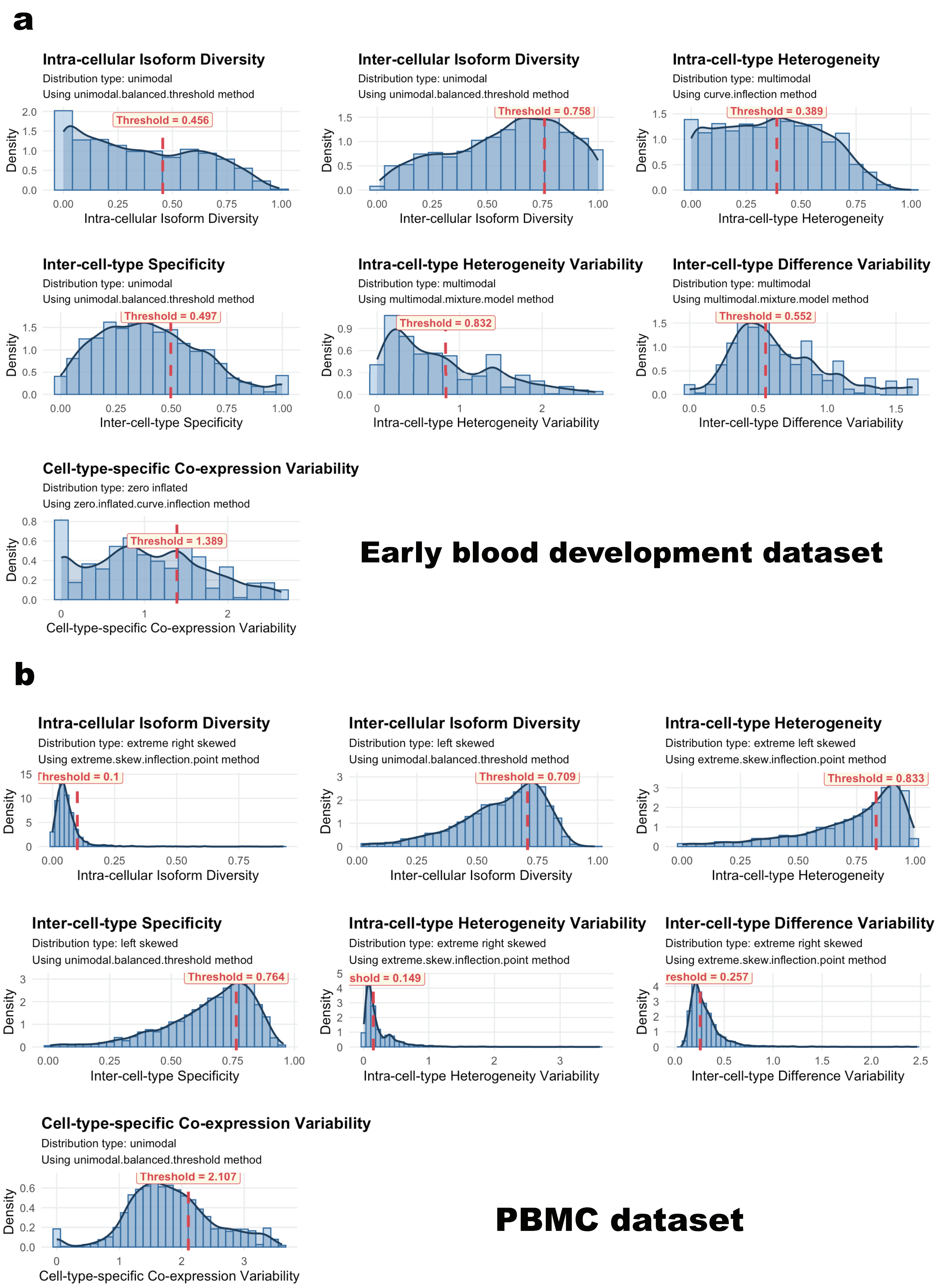


Supplementary Figure 7: **Visualisation of complexity metrics threshold determination for early blood development and PBMC datasets.**

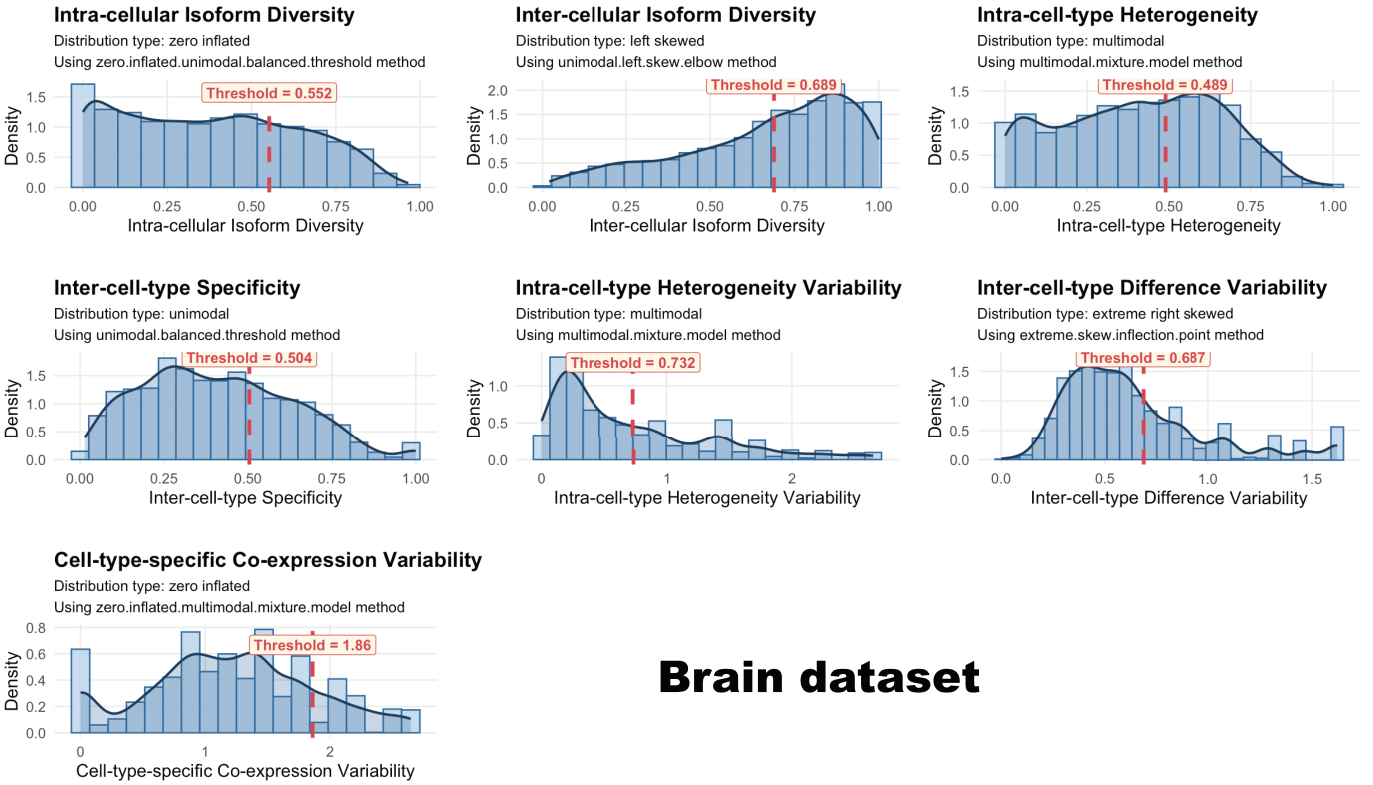


Supplementary Figure 8: **Visualisation of complexity metrics threshold determination for brain dataset.**
